# Tracing the evolutionary history of Tunisian HPV 16 & 18 across the genetic diversity in L1 gene

**DOI:** 10.1101/2024.02.15.580472

**Authors:** Monia Ardhaoui, Naira Dekhil, Zied Bouslama, Rahima Bel Hadj Rhouma, Emna Fehri, Emna Ennaifer, Ikram Guizani

**Affiliations:** Molecular Epidemiology and Experimental pathology-LR16IPT04, Pasteur Institute, Tunis, University of Tunis El Manar, Tunis, Tunisia; Laboratory of Anatomic pathology, Pasteur Institute, Tunis, University of Tunis El Manar, Tunis, Tunisia; Unit of Typing & Genetics of Mycobacteria, Laboratory of Molecular Microbiology, Vaccinology, and Biotechnology Development-LR16IPT01, Pasteur Institute, Tunis, University of Tunis El Manar, Tunis, Tunisia; Laboratory of Epidemiology and Veterinary Microbiology - LR16IPT03, Pasteur Institute, Tunis, University of Tunis El Manar, Tunis, Tunisia

## Abstract

HPV16 and HPV18 are the most prevalent high risk HPV types, recognized to be HPV vaccine target. Currently, available vaccines against HPV infections are based on virus-like particles (VLPs) derived from the L1 protein. Understanding the emergence of HPV isolates and exploring the genetic diversity of their L1 gene is crucial to assess the effectiveness of vaccine introduction in different populations. This is the first study aiming to investigate the evolutionary dynamics of HPV16 and HPV18 variants circulating in Tunisia. We constructed evolutionary history of HPV16 and HPV18 based on partial L1 sequence dataset of 733 Tunisian and worldwide representative isolates. Phylogeographic analysis confirmed the European origin of Tunisian strains that have emerged in 90’s. Strikingly, four nonsense mutations in HPV16 and one in HPV18 within the HI surface loop, a critical immunogenic region were identified. Moreover, the latter showed an excess of non-synonymous over synonymous substitutions, a hallmark of local adaptation. By elucidating the genetic variability in the L1 gene, our study unveils several new features that pose challenges to the ongoing fight against HPV. These insights underscore the importance of continuous surveillance and adaptation of vaccination strategies to address evolving viral dynamics effectively.

**Importance:** Human Papillomavirus (HPV) is a virus known to cause cervical cancer (CC). HPV16 and HPV18 are the most common HPV types related to CC. This study aimed to trace the evolutionary history of Tunisian HPV16 and HPV18 isolates. Phylogeographic analysis showed that these viruses were originated from Europe and emerged in 1990’s. Several genetic changes have been identified in the L1 gene of HPV16 & 18 that could potentially reduce the effectiveness of HPV vaccines. By understanding these changes and addressing the local adaptation of HPVs, we could prevent the spread of this virus within a population.

## Introduction

Human papillomaviruses (HPVs) are double-stranded DNA viruses that infect mucosal and cutaneous epithelia, contributing to various cancers, notably cervical cancer (CC) [1-3]. The classification of HPVs is based on the sequence similarity of the L1 gene, leading to five genera: Alphapapillomaviruses (alpha), Betapapillomaviruses (beta), Gammapapillomaviruses (gamma), Mupapillomaviruses (mu), and Nupapillomaviruses (nu) [4, 5]. HPV genomes are further categorized into intratypic variants based on their genome sequence diversity, with lineages exhibiting a nucleotide divergence of 1% to 10% and sub-lineages displaying a divergence of 0.5% to 1% [6-9]. Mucosal alpha-PVs, designated as high-risk (HR-HPV) and low-risk (LR-HPV), are associated with carcinogenic potential [4, 5]. Notably, 15 HPV genotypes are classified as high-risk (e.g., HPV16, 18, 31) and 12 as low-risk (e.g., HPV6, 11) [10]. Persistent HR-HPV infection, particularly with HPV16 and HPV18, contributes significantly to cervical cancer development, representing 70% of CC worldwide [11].

Previous studies on HPV16 and HPV18 variants have identified three lineages based on continental locations of viral samples —European (E), Asian-American (AA), and African (Af) [12]. The phylogeny of these variants reflects human migration patterns, suggesting divergence through genetic isolation concurrent with Homo sapiens’ establishment in different continental regions. While direct detection of HPV DNA in archaeological remains is challenging, co-migration with modern human populations after the out-of-Africa migration likely influenced the global distribution of HPV genotypes [2, 3].

In Tunisia, HPV16 prevails as the most common high-risk type, with prevalence rates ranging from 16% in women with normal cervical cytology to 60% in those with cervical intraepithelial neoplasia or cervical cancer [13-17]. Although HPV18 is less frequently detected than HPV16, its significance persists, commonly identified in cervical cancer and targeted by vaccines.

Understanding the dynamic evolution of HPV16 and HPV18 is crucial for developing effective prevention and vaccination programs. However, in Tunisia, the emergence and spread of these viruses are little explored. This study aimed to trace temporal and geographical origin of HPV16 and HPV18 isolated from Tunisian women. It contributes to the broader understanding of the genetic diversity, transmission patterns, and evolutionary history of the virus in North Africa.

## Methodology

### Data collection

Data for this study involved Tunisian sequences, references and publically available L1 gene sequences (N= 733) as described below.

References sequences (n=24) from different HPV16 and HPV18 lineages were obtained from the Pave Database (https://pave.niaid.nih.gov).

Tunisian L1 gene of HPV 16 (n =32) and HPV18 (n = 29) were generated from a previous Tunisian national epidemiological study that aimed to investigate the prevalence and distribution of human papillomavirus (HPV) genotypes in the Tunisian population [18]. The sequences were submitted to Genbank under the accession number PP212995-PP21302 and PP327891-PP327919 for HPV16 and HPV18 respectively.

Additionally, complete or partial L1 gene sequences were retrieved from NCBI virus (https://www.ncbi.nlm.nih.gov/labs/virus/vssi/) databases. Only sequences with the collection date and geographic data were kept for further analysis. Overall, n=371 and n=301 sequences for L1-HPV16 and L1-HPV18 respectively were retained (Supplementary data S1).

### Phylogenetic analysis

For the identification of HPV16 and HPV18 Tunisian lineages based on L1 sequences, different linages reference and Tunisian sequences were aligned using the ClustalW program in MEGA11 software [19]. Maximum-likelihood (ML) trees under GTR+T4 model were constructed using. The best tree was selected after 1000 bootstraps.

### Divergence times and evolution analysis

Divergence times and evolutionary rates were calculated using LSD2 (Least Squares Dating), a molecular clock method based on least-squares optimization [20]. Phylogenies were generated using Iqtree [21] with 1000 bootstraps and selecting the best-scoring X74477_HPV35 genome was used as outgroup. To overcome the limitation of testing a single tree topology and a strict clock, we performed a date randomization test with 1000 randomized datasets using the quadratic programming dating (QPD) algorithm and calculated the confidence interval (options -f 1000 and -s). The tree was visualized and annotated using iTOL version 5 [22].

### Phylogeographic analysis

Phylogeographic analysis was performed using PASTML software, which is based on different models and inference methods [23]. To investigate the origin of Tunisian HPV clinical strains in a worldwide context, we used a globally distributed L1 gene dataset consisting of 733 isolates covering a wide temporal and geographic range. Marginal posterior probability approximation (MPPA) was applied with an F81-like model.

### Genetic diversity of HPV 16 & 18 in L1 gene

For genetic diversity analysis, partial L1 gene sequences from HPV16 and HPV18 (142 bp, Genomic position: 5762-5903) were aligned with reference genomes using MEGA11 software. Alignment visualization was performed using the Weblogo tool [24]. Non-synonymous Single Nucleotide Polymorphic sites (nsSNPs) were mapped to HI loop region using SwissModel [25].

dn/dS ratio was then calculated using Dnasp software [26] to assess selective pressures.

## Results

Phylogenetic based on partial L1 gene (N=32 sequences) showed that 37% of Tunisian HPV16 variants belonged to European lineage A, followed by 34% to American and Asian lineages (C/D lineages), and 25% to African lineage (B lineage). One sample didn’t belong to any lineage (Figure 1). Phylogenetic tree of HPV18 L1 gene sequences (N=29) revealed that all Tunisian isolates belonged exclusively to lineage A (19/29, 90.5%) (Figure 2).

**Figure 1.**
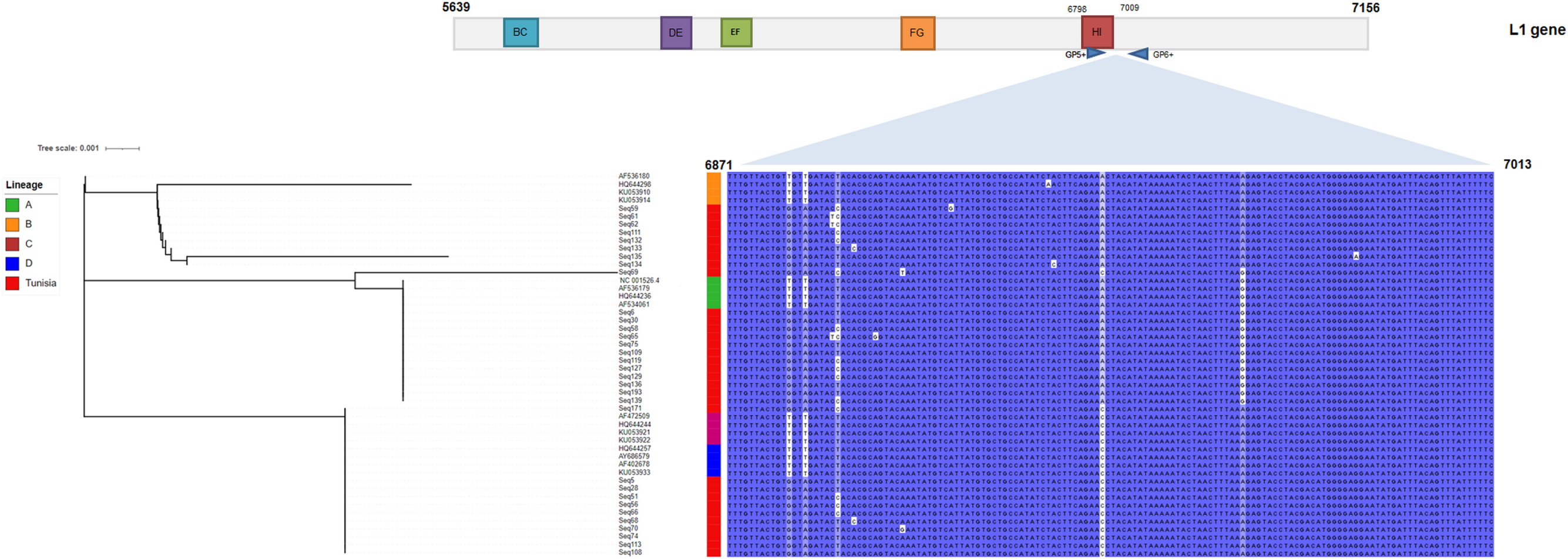
Maximum likelihood phylogenetic tree based on partial L1 gene of Tunisian HPV16 isolates and their relative sequences used in this study. Seq represents Tunisian sequences, NC 001526, AF536179 (A2), HQ644236 (A3), AF534061 (A4), AF472509(C1), HQ644244 (C2), KU053921 (C3), KU05922 (C4), HQ644257 (D1), AY68579 (D2), AF4022678 (D3), AF531680 (B1), KU053914 (B4), AF536180 (B1), HQ644298 (B2), KU053910 (B3) correspond to reference sequences from the different lineages.

**Figure 2.**
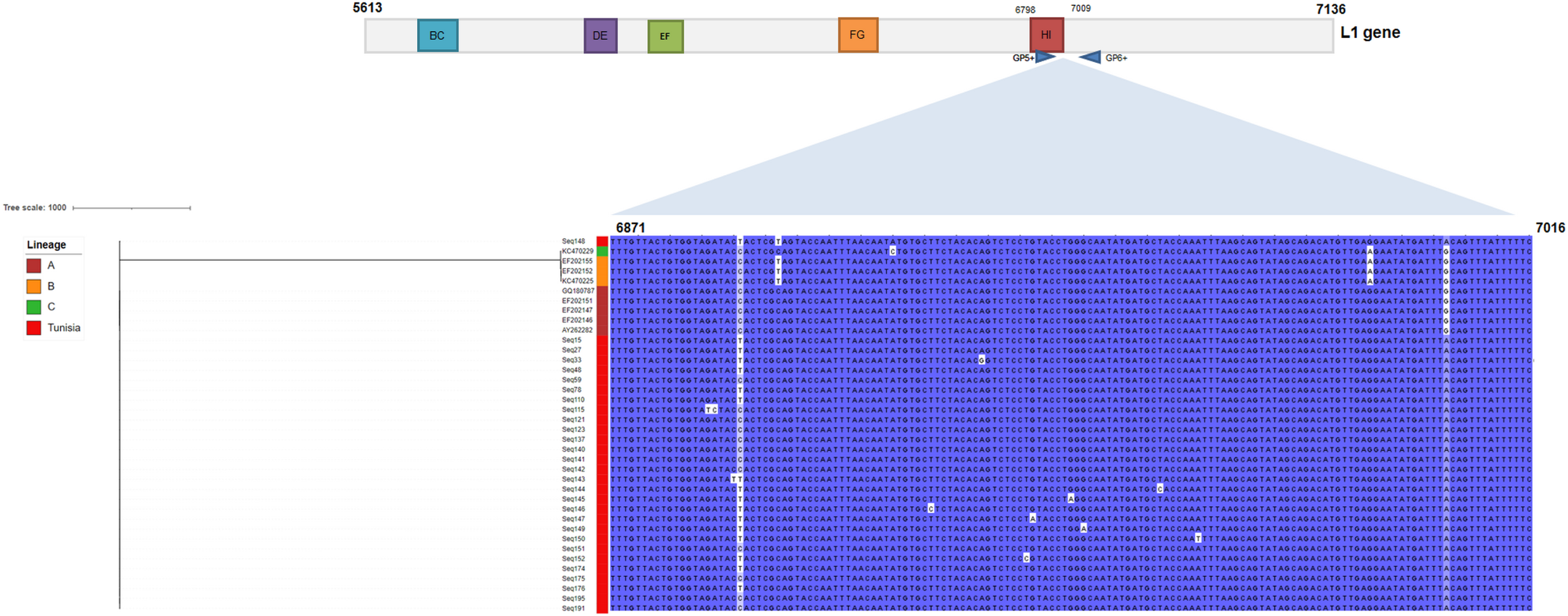
Maximum likelihood phylogenetic tree based on partial L1 gene of Tunisian HPV18 isolates and their relative sequences used in this study. Seq represents Tunisian sequences. AY262282 represents reference sequences of HPV18 (Lineage A1), EF202146 (A2), EF202147 (A3), EF202151 (A4), GQ180787 (A5), EF202155 (B1), KC470225 (B2), EF202152 (B3), KC470229 (C1).

### Phylogeographical analysis of Tunisian HPV16/HPV18 L1 gene

Phylogeographic analysis was performed using PASTML on 371 and 301 sequences of HPV16 and HPV18 respectively. HPV16 was mainly derived from Italy cluster (Figure 3) and HPV18 from Netherlands cluster (Figure 4). Strikingly, both HPV seemed to be emerged from the same cluster as European isolates.

**Figure 3:**
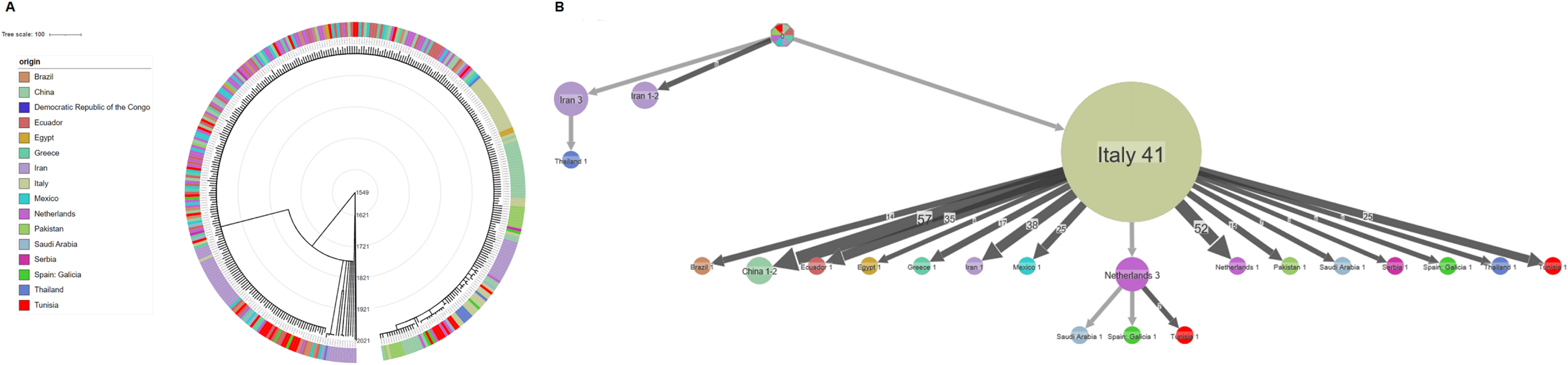
Spatio-temporal analysis of HPV16. **A**: Time-scaled phylogenetic tree HPV16 isolates based on partial L1 gene. External colour strips indicate the origin of clinical isolates. Temporal predictions are obtained with LSD2 and the visualisation is performed with iTOL version 5. **B**: Dispersal of Tunisian HPV16 based on partial L1 gene. Different colours correspond to different geographical regions. Numbers inside the circles indicate the number of strains assigned to the specific node. External color strips indicate the origin of clinical isolates. Temporal predictions are obtained with LSD2 and the visualization is performed with iTOL version 5.

**Figure 4:**
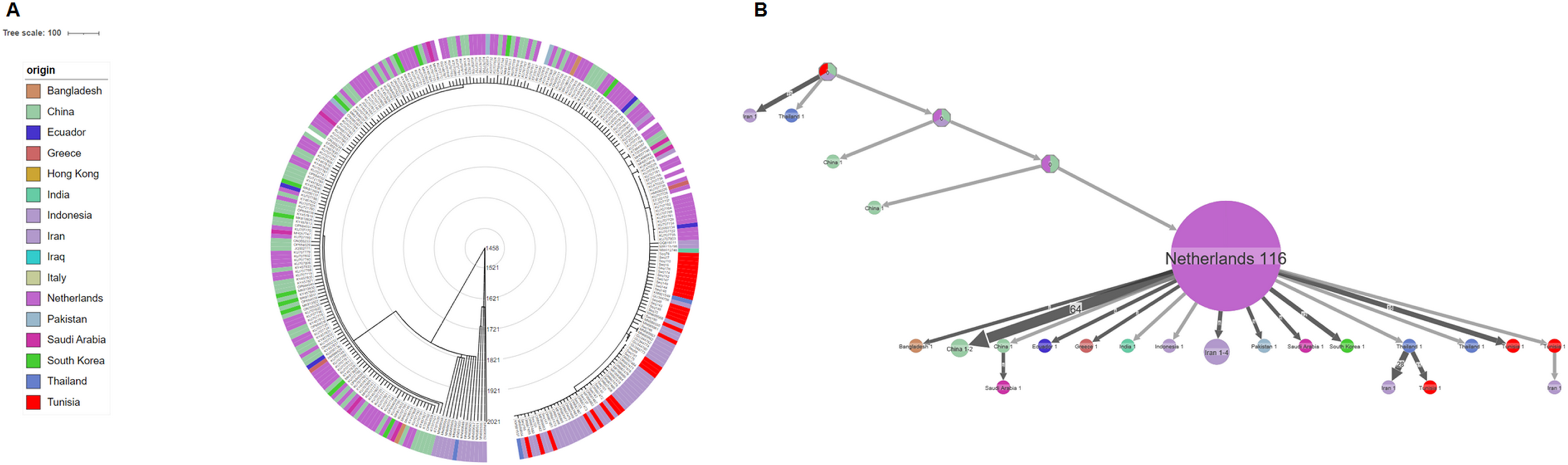
Spatio-temporal analysis of HPV18. **A**: Time-scaled phylogenetic tree HPV18 isolates based on partial L1 gene. External colour strips indicate the origin of clinical isolates. Temporal predictions are obtained with LSD2 and the visualisation is performed with iTOL version 5. **B**. Dispersal of Tunisian HPV18 based on partial L1 gene. Different colours correspond to different geographical regions. Numbers inside the circles indicate the number of strains assigned to the specific node. The colour code for origin is following the legend 3. Circles are proportional to the number of isolates, and the arrows are proportional to the number of times the migration has been observed on the tree. The branches represent the inferred ancestral region with the largest posterior probability.

### Time of the most recent common ancestral of Tunisian HPV16 & 18 isolates

Table 1 corresponds to the mutation rate per site/year and time to the most recent common ancestor (TMRCA) based on partial L1 gene of HPV16 and HPV18. Results showed that HPV16 emerged in 1991 95% CI [1983-2003], slightly before HPV18 1993 95% CI [1980-1998] in Tunisia. However, HPV18 displayed higher mutation rate than HPV16.

**Table 1.** Divergence time estimates Tunisian HPV16 and HPV18 isolates MRCAs.

|  | **Mutation rate [95% CI]** | **TMRCA [95% CI]** |
| --- | --- | --- |
| Overall HPV18 | 2.0e-05 [1.16e-05; 2.59e-05] | 1457 [1029.35; 1614.02] |
| Tunisian HPV18 | 2.0e-05 [1.16e-05; 2.59e-05] | 1993.86 [1980.26-1998.32] |
| Overall HPV16 | 5e-07 [5e-07; 1.2e-06] | 1548.9 [1157.85; 1871.45] |
| Tunisian HPV16 | 5e-07 [5e-07; 1.2e-06] | 1991.01 [1983.72;2003.24] |

### Genetic diversity of HPV16 & 18 isolates circulating in Tunisia

Genetic diversity analysis within partial HPV16 L1 gene sequences from Tunisian samples unveiled four non-synonymous nucleotide changes: T362I (3/32), S365G (1/32), T376P (1/32), and T379P (12/32). Similarly, examination of partial HPV18 partial L1 gene sequences revealed five non-synonymous nucleotide polymorphisms: D396S (1/29), T397I (1/92), Q410R (1/29), G415R (1/29), and K421N (1/29). Using SwissModel software, the identified nsSNPs within the partial L1 sequences of HPV16 and HPV18 were found to be situated within the HI loop region (Figure 5). Details of these findings, including dn/ds ratios, are presented in Table 2.

**Figure 5.**
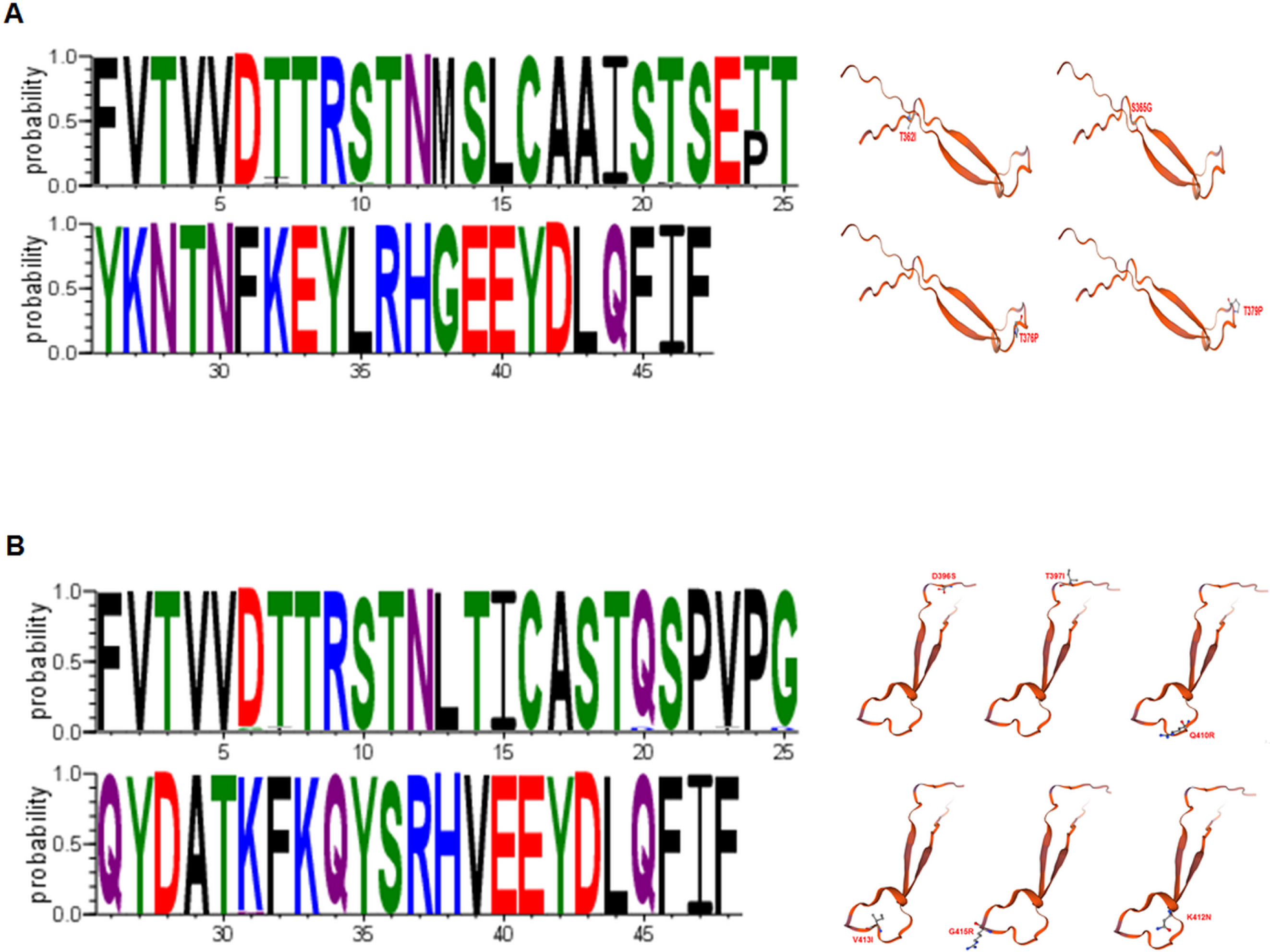
A. Weblog representation of amino acids variation within partial HPV16 and HPV18 L1 protein. B. nsSNP maps in partial HPV16 and HPV18 L1 HI loop region.

**Table 2.**
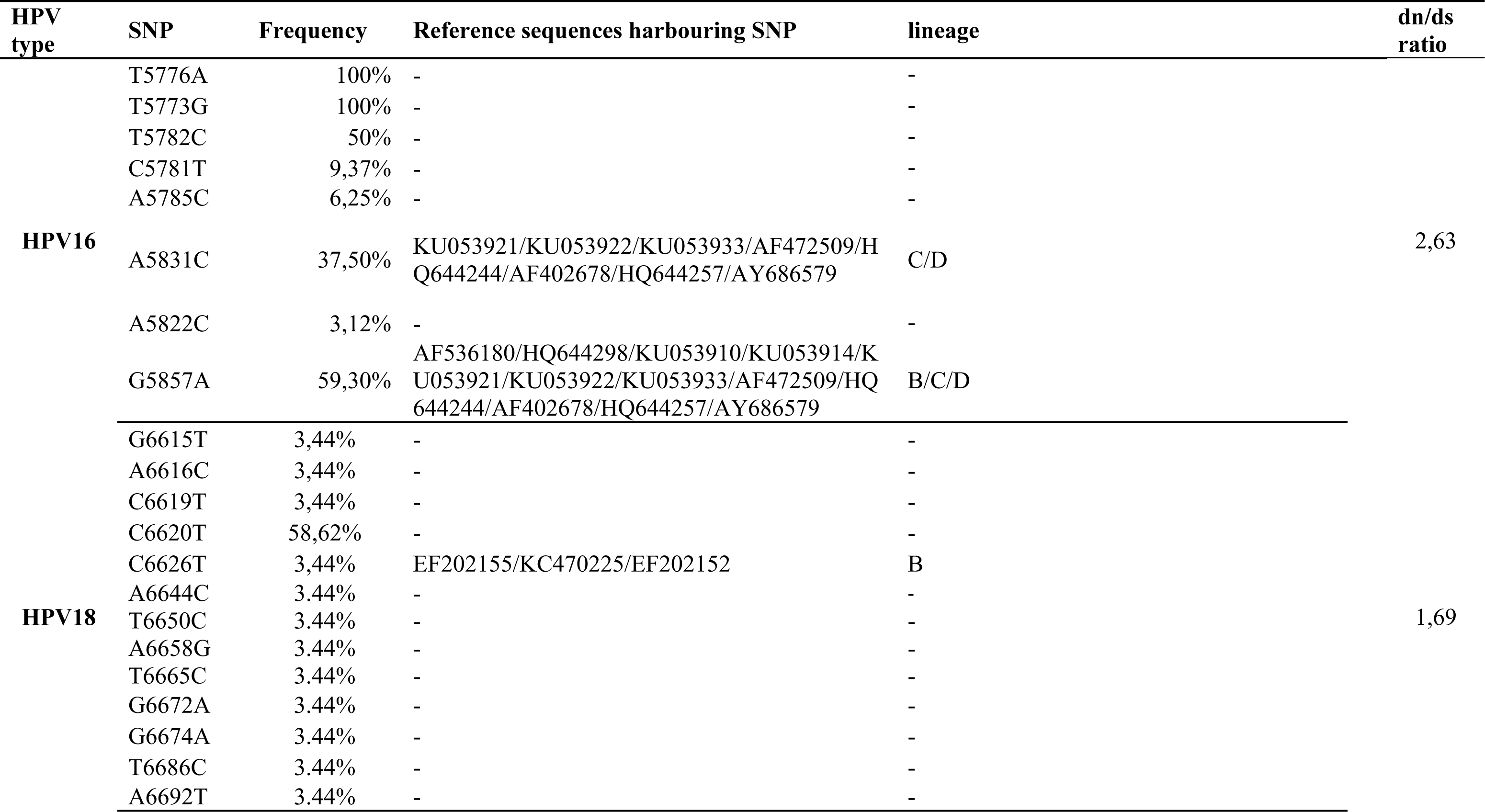
Summary of identified mutations in the partial L1gene sequences and dn/ds ratio in HPV16 & 18.

## Discussion

L1 gene has been the focus of many studies, as it encodes the major capsid protein L1 and its potential impact on the vaccination-induced immunity [27-29]. HPV 16 and HPV 18 are the most prevalent infection types among cervical cancer patients, responsible for ∼71% of cervical cancers [30]. Here, we conducted genetic diversity and evolutionary dynamics analysis of Tunisian HPV 16 and HPV18 isolates based on a partial L1 sequence. Results highlighted the genetic closely related HPV16 and HPV18 to European Lineage A, with respectively 37% and 100%. Taking into account the high prevalence of European HPV Lineage in Mediterranean Women [31-34], our finding confirmed its dispersal among Tunisian Women. Moreover, the lineage A is dominant in Algeria and Morocco, but lineages B and C are mainly detected in Sub-Saharan Africa [33]. This predominance is explained by the history of European migration to North Africa.

Previous study has reported that HPV16 emerged in Croatia in 1987, then spread to Italy in 1994 and five years later (1999) to Spain [35]. The latter, has prompted us to conduct Bayesian analysis to elucidate the emergence of HPV Tunisian isolates. The mutation rate based on L1 partial sequence was estimated to 5 e-7 per site per year for HPV16. This rate is similar to the upstream regulatory region (URR) (∼4.5 e−7 per site per year) [36] and aligns with molecular clock studies, which propose a typically slow evolution of HPV, estimated at approximately 1% nucleotide change per 100,000 to 1,000,000 years [37]. Surprisingly, this rate seemed to be higher in HPV18. This result could be explained by the fact that much more SNPs have been detected in the partial L1 sequence of HPV18 despite their low frequency.

The TMRCA estimates based on L1 partial sequence in HPV18 and HPV16 suggest a modern introduction of these variants in Tunisia (90’s). Interestingly, the 1994 date is consistent with the reported emergence of HPV in Italy [35]. Furthermore, phylogeographic analysis confirmed the European origin of Tunisian HPV, as they derived from the same cluster as Italian and Netherland (Figure 3B and Figure 4B). Thus, it marks the migration of HPV from Europe to North Africa. This observation lends support to the notion that the topologies of HPV16 and HPV18 phylogenies may indeed mirror the history of modern human migration [17, 18].

L1 partial sequences used in this study covered 17% of HI loop, one of the five hypervariable surface loops in this gene (BC, DE, EF, FG, and HI), and known to be targets for recognition by human antibodies during immune responses [38, 39]. Four non synonymous SNPs have been detected in HPV 16 (T362I, S365G, T376P, T379P) and one in HPV18 (D396S) (Figure 5). Notably, mutations occurring in surface loops, or in sites between them, have been reported to potentially impact virus infection efficiency or alter virus antigenicity [28]. In silico predictions suggested that T379P may lead to a 0.4-fold decrease in antibody titer, highlighting the impact of SNPs in HI loop on L1 epitopes and immunogenicity [40]. Moreover, genetic diversity analysis showed that this region was under positive selection, revealed by a dn/ds ratio of 2.63 and 1.69 in HPV16 and HPV18. Our data highlight genetic variability that could affect the vaccination-induced immunity on Tunisian population and the selective pressure that vaccinations apply to high-risk HPV strains, leading to competition between strains that are trying to survive in the niche. Further investigation for the whole genome is needed to well-characterize HPV variants and understand their adaptation capacity within a population.

## Conclusion

The present study provides initial insights into the evolutionary history and genetic diversity of HPV16 and HPV18 based on partial L1 gene. Our analyses reveal that the ancestral strains of Tunisian HPV16 and HPV18 can be traced back to 90’s, with a potential origin from Europe. This study represents a pioneering effort in investigating the emergence and adaptation of HPV in Tunisia. Such exploration contributes valuable perspectives to our comprehension of the dynamic evolutionary processes within a region characterized by historical interactions through colonization and immigration.

## Supplementary data

**Supplementary data S1.** HPV16 & 18 Metadata: Origin and date of published sequences of gene L1 of HPV16 and HPV 18

